# Intrinsically disordered interaction network in an RNA chaperone revealed by native mass spectrometry

**DOI:** 10.1101/2022.05.23.493136

**Authors:** Samantha H. Sarni, Jorjethe Roca, Chen Du, Mengxuan Jia, Hantian Li, Ana Damjanovic, Ewelina M. Małecka, Vicki H. Wysocki, Sarah A. Woodson

## Abstract

RNA-binding proteins contain intrinsically disordered regions whose functions in RNA recognition are poorly understood. The RNA chaperone Hfq is a homohexamer that contains six flexible C-terminal domains (CTDs). The effect of the CTDs on Hfq’s integrity and RNA binding has been challenging to study because of their sequence identity and inherent disorder. We used native mass spectrometry (nMS) coupled with surface-induced dissociation (SID) and molecular dynamics (MD) simulations to disentangle the arrangement of the CTDs and their impact on the stability of *E. coli* Hfq with and without RNA. The results show that the CTDs stabilize the Hfq hexamer through multiple interactions with the core and between CTDs. RNA binding perturbs this network of CTD interactions, destabilizing the Hfq ring. This destabilization is partially compensated by binding of RNAs that contact multiple surfaces of Hfq. By contrast, binding of short RNAs that only contact one or two subunits results in net destabilization of the complex. Together, the results show that a network of intrinsically disordered interactions integrate RNA contacts with the six subunits of Hfq. We propose that this CTD network raises the selectivity of RNA binding.

**Significance Statement:** Hfq is a protein hexamer necessary for gene regulation by non-coding RNA in bacteria, during infection or under stress. In the cell, Hfq must distinguish its RNA partners from many similar nucleic acids. Mass spectrometry dissociation patterns, together with molecular dynamics simulations, showed that flexible extensions of each Hfq subunit form a dense network that interconnects the entire hexamer. This network is disrupted by RNA binding, but the lost interactions are compensated by RNAs that contact multiple Hfq subunits. By measuring interactions that are too irregular to be counted by other methods, mass spectrometry shows how flexible protein extensions help chaperones like Hfq recognize their RNA partners in the messy interior of the cell.

## Introduction

Many RNA-binding proteins (RBPs) contain intrinsically disordered regions (IDRs) (1) with overlapping functions that have been difficult to disentangle. For example, IDRs may augment specific RNA recognition, connect different RNA-binding modules, and enable the assembly of liquid condensates, while also serving as targets for post-translational modification (2–4). The heterogeneous and dynamic structures of IDRs make their interactions especially challenging to quantify, and their functions in most RNA binding proteins remain poorly understood.

Hfq is a bacterial Sm protein that binds small non-coding RNA (sRNA) and chaperones sRNA regulation of complementary mRNAs (5) (Fig. 1). Deletion of Hfq results in pleiotropic effects, including maladaptive responses to stress and decreased virulence (6). The well-folded core of the Hfq hexamer assembles into a symmetric ring that binds U and A-rich sequence motifs in sRNA and mRNA substrates (7). Conserved arginine patches on the outer rim of the hexamer also bind RNA and are essential for its chaperone activity (8, 9).

**Fig. 1.**
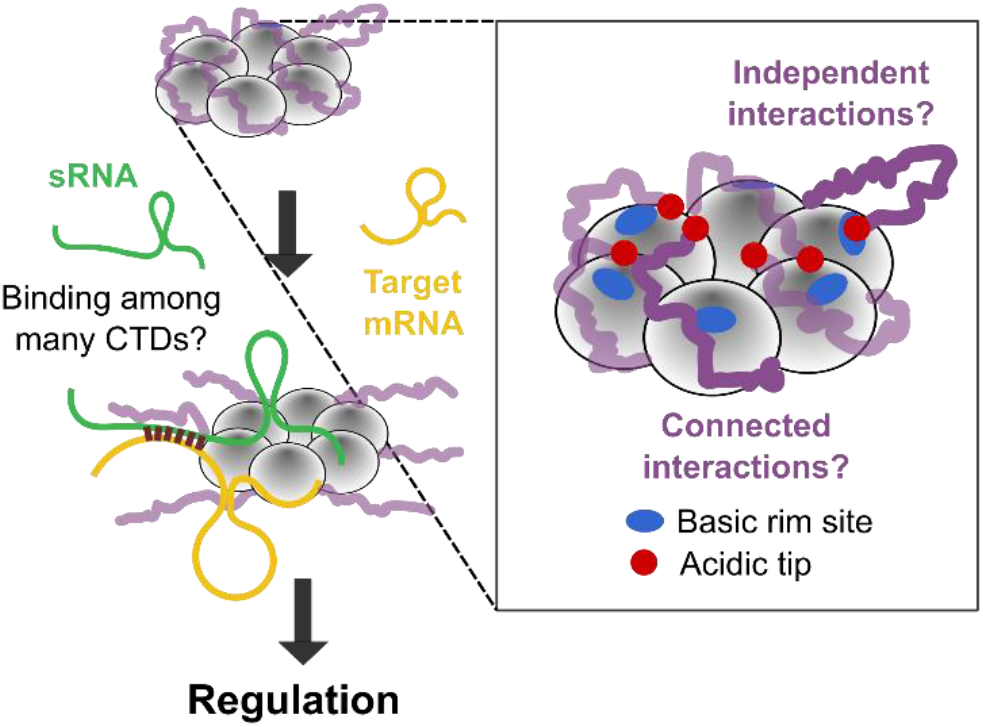
Role of Hfq’s CTDs in sRNA regulation. Hfq chaperones the annealing of sRNAs with their target mRNAs, but it is not known how binding of RNAs occurs when the core of Hfq is occluded by many disordered CTDs. Although the acidic tips of the CTDs (red) can interact with basic patches on the rim (blue) (21), the organization and collective behavior of the CTDs is unknown.

*E. coli* Hfq also has intrinsically disordered C-terminal domains (CTDs) that extend outward from the core of the hexamer (10). Each monomer containing 102 residues contributes a 37 amino acid (aa) CTD, creating a crowded zone of disordered polypeptide around the protein. This ring-shaped organization, which is unlike disordered regions in other RBPs, raises the possibility that the Hfq CTDs act together rather than individually.

Despite their disordered nature, NMR chemical shift perturbations and molecular dynamics simulations determined that the CTDs of *E. coli* Hfq interact with the rim of the hexamer (11). Additionally, unassigned electron density in a crystal structure of Hfq bound to RydC sRNA suggested that the CTDs make distributed contacts with the protein-RNA surface (12). These results aligned with the earlier observation that the CTDs (residues 65-102) stabilize the Hfq hexamer (13) and contribute to its function (14–19). We found that semi-conserved acidic residues at the C-terminus mimic nucleic acid, competing with RNA for binding to the rim (20–22). Competition with the CTDs can result in preferential dissociation of non-specific RNA, and retention of specific RNA ligands. More recently, it was shown that the bases and the tips of the CTDs interact synergistically with particular Hfq surfaces, leading to different effects depending on the RNA ligand (23).

Because of their intrinsic disorder and sixfold symmetry, how the CTDs organize around Hfq stabilizing the hexamer is still unknown. Additionally, it is not known if each CTD acts locally and independently, or if the six CTDs act together to accommodate or displace an incoming RNA (Fig. 1). Moreover, the energetic contributions of individual CTDs to RNA binding have been almost impossible to quantify.

We addressed these challenges by using native mass spectrometry (nMS) coupled with surface-induced dissociation (SID) (Figs. 2*A* and S1). In nMS, the protein complex is exchanged into a volatile electrolyte allowing transfer of the intact native complex to the gas phase (24). After ionization, collision with a surface (nMS-SID) dissociates the complex into product ions that provide information about the stabilities of the non-covalent interfaces within the complex and their molecular organization (25). This method has been used to characterize the stability, structure and assembly pathways of many protein complexes, including RBPs and membrane proteins (26–28). Yet, despite its promise for discovery, nMS-SID studies of large biomolecular complexes typically require customized instrumentation (Fig. S1).

**Fig. 2.**
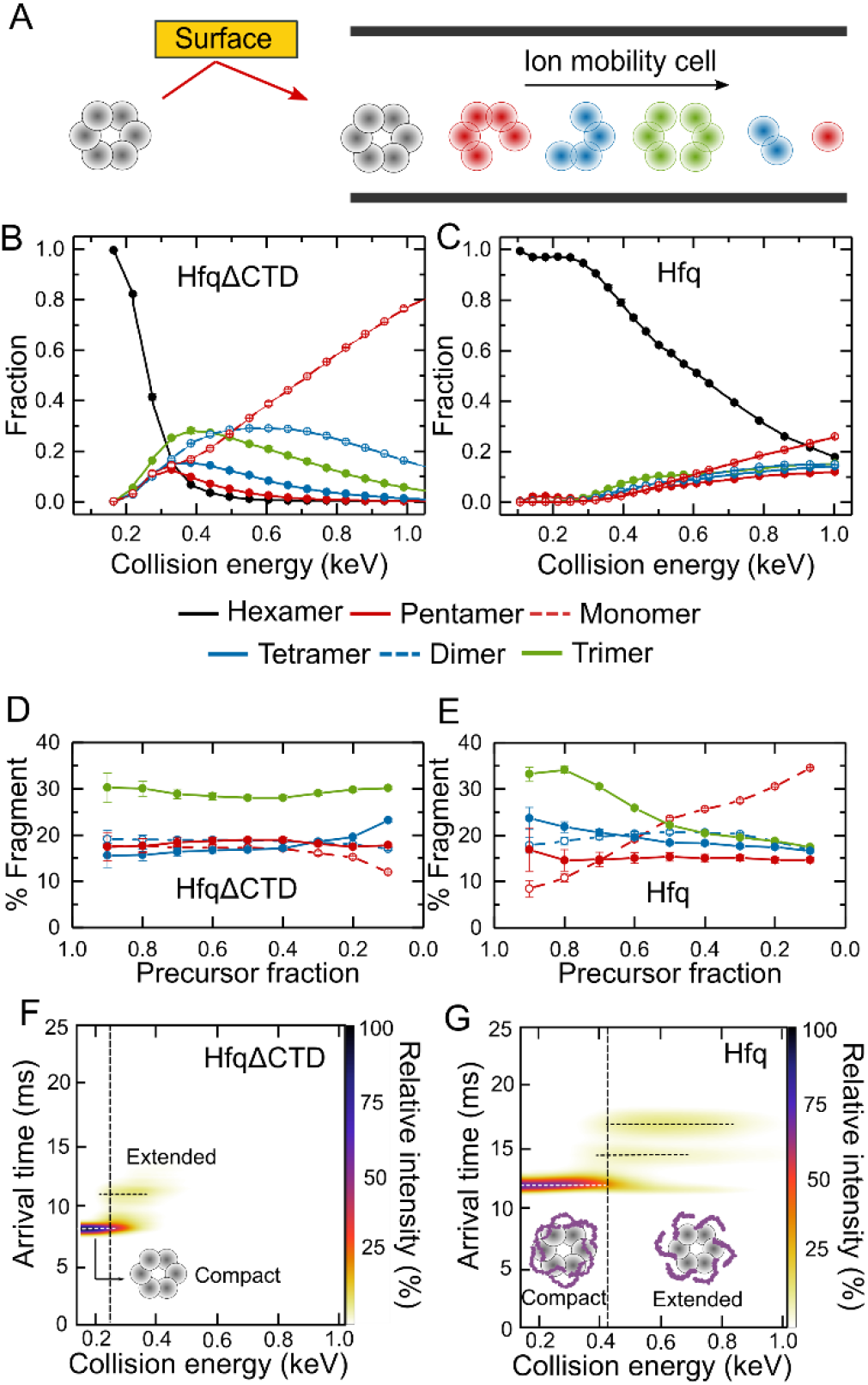
Disordered CTDs stabilize Hfq. (*A*) Native mass spectrometry coupled with surface-induced dissociation (nMS-SID) dissociates the Hfq hexamer into oligomers that retain the connectivity of the native protein. Fragments are separated according to their arrival time after traversing an ion mobility cell. (*B-C*) Energy-resolved mass spectrum of (*B*) HfqΔCTD and (*C*) Hfq. The collision energies are corrected for the mass of the CTDs (m_HfqΔCTD_/m_Hfq_). Reported fractions are the sum of the intensities of each dissociation product normalized by the total intensity of all products. Symbols report the average of three replicates. Standard errors are smaller than the symbols for some data points. Solid lines represent a cubic interpolation of the data. (*D-E*) Percentage of each oligomer (pentamer, tetramer, trimer, dimer or monomer) in the dissociation products, as a function of the remaining precursor fraction for (*D*) HfqΔCTD and (*E*) Hfq. Errors are the spread of the ERMS curves, normalized by the total dissociated fraction and converted to a percentage. Colored as in Figs. 2*B-C*. Solid lines are a visual guide. (*F-G*) Surface-induced unfolding (SIU) plot of (*F*) HfqΔCTD and (*G*) Hfq. The dashed line represents the transition from compact to extended protein and corresponds to ~ 0.2 and ~ 0.7 precursor fraction for each protein, respectively.

Here, by using nMS-SID and all-atom molecular dynamics (MD) simulations, we show that the six disordered CTDs of apo Hfq form extensive interactions that energetically connect and stabilize the entire hexamer. When RNA binds any subunit of Hfq, these stabilizing interactions are disrupted throughout the hexamer. Taken together, our results show how disordered regions can integrate RNA-protein interactions across a multi-subunit chaperone.

## Results

### Disordered CTDs stabilize Hfq

To better understand the interactions of the flexible CTDs, we first used nMS-SID on WT *E. coli* Hfq (102 aa per subunit) and a truncated Hfq lacking the CTDs (HfqΔCTD; 65 aa per subunit). We analyzed the dissociation products (pentamer, tetramer, trimer, dimer, and monomer) obtained at increasing collision energies (CE) with the surface (Fig. 2*A*). The resulting energy-resolved mass spectra (ERMS) showed that the two proteins have different stabilities and fragmentation patterns (Figs. 2*B-C*). HfqΔCTD reached 20% fragmentation (80% precursor remaining) at ~220 eV, compared to ~390 eV for WT Hfq (Figs. 2*B-C*, black lines). This large difference demonstrated that the wild-type protein is much more stable than the truncated version, in agreement with previous reports (13).

The fraction of HfqΔCTD hexamer sharply decreased with modest increases in collision energy, with 90% fragmentation of the precursor at CE ~350 eV. In contrast, the amount of WT Hfq dissociation increased gradually over a wide range of collision energies, and ~20% of the precursor remained intact even at CE = 1000 eV. This gradual response to higher collision energies suggested that the CTDs prevent dissociation of the complex across a wide range of energies. One explanation is that the CTDs form a range of intersubunit interactions that reorganize upon activation. Additionally, the CTDs may be organized differently in each Hfq hexamer, causing dissociation over a continuum of collision energies.

### Disordered CTDs impact the connectivity of Hfq

The dissociation patterns for HfqΔCTD and Hfq generated fragments in different ratios, indicating a different degree of connectivity between subunits in the two proteins (Figs. 2*B-E*). We compared the dissociation products of the two proteins at energies that deplete the hexamer precursor equally. HfqΔCTD dissociated into twice as many trimers (~30%), as other fragments (~15–20%) regardless of the total amount of precursor dissociated (Fig. 2*D*). This result suggested that the protein interfaces around the ring have a similar probability of dissociating.

For WT Hfq, the distribution of dissociation products was markedly different from HfqΔCTD (Fig. 2*E*), indicating that the CTDs contribute to the subunit interfaces, as also reported earlier (13, 29). For all fractions of the precursor, the percentage of pentamers, tetramers and dimers remained similar (~10-20%). However, the percentage of trimers decreased while the percentage of monomers increased as more precursor was fragmented.

Inspection of ion mobility arrival times revealed that as the collision energy increases, the WT hexamer precursor converts into two complexes that migrate more slowly in the drift chamber, suggesting partial extension or restructuring of the protein hexamer, although we can’t conclude that this occurs before the subunits dissociate from one another (surface-induced unfolding (SIU) plots, Figs. 2*F-G*). We observed only one additional complex of HfqΔCTD upon activation (Fig. 2*F*), suggesting that one of the extended Hfq complexes comes from restructuring of the core beta sheet. The second extended form was only observed for WT Hfq and likely arises from extension of the CTDs (Fig. 2*G*). Altogether, the nMS results support a model in which the CTDs are bound to the core and each other in WT Hfq, stabilizing the entire hexamer. At increasing collision energies, the CTDs disentangle, exposing the core and eliminating the stabilizing inter-subunit connections. As a result, the fragile hexamer ruptures into smaller complexes.

### MD simulations reveal a network of CTD interactions on Hfq

To gain more insights into the organization of the disordered CTDs on Hfq, we performed multiple all-atom molecular dynamics (MD) simulations on the wild-type protein. The simulations were started from 10 previous low-energy Rosetta structures (models top 1-10) (21) and an extended structure started with 4 different initial velocities (extended models 1-4). The structures obtained after equilibration agreed with the experimentally determined collisional cross section (CCS), with the MD structures being more compact than the initial Rosetta models (Fig. S2 and Table S1). The simulated CTDs adopted a variety of conformations (Figs. 3*A* and S3). Thus, the diverse organization of the disordered CTDs confers heterogeneity to individual Hfq hexamers, as implied by the ERMS results (Fig. 2*C*).

**Fig. 3.**
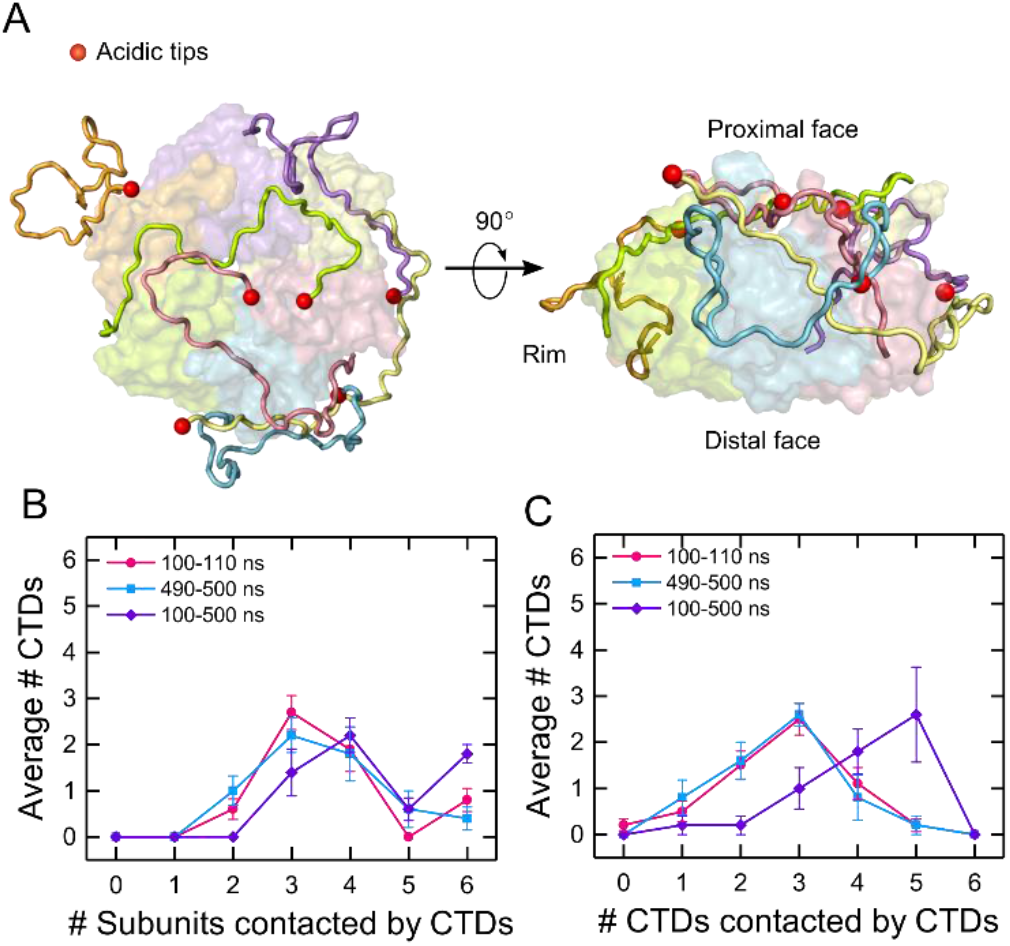
Disordered CTDs create a network of interactions with Hfq. (*A*) (*Left*) Top and (*right*) side view of an Hfq model from one molecular dynamics (MD) simulations (top 4, at 500 ns). Subunits are colored individually with each acidic C-terminus shown as a red sphere. (*B-C*) Average number of CTDs interacting with 0, 1, 2, 3, 4, 5 or 6 (*B*) subunits or (*C*) CTDs, for all MD models. Interactions were considered at any time during the first 100-110 ns for all models, the last 490-500 ns for models top 1-5 and for the entire simulation (100-500 ns) for models top 1-5. Symbols are the average and standard error of the mean (SEM) of the various simulations. Solid lines are a visual guide.

The Hfq models showed that each CTD may interact within its monomer and with other subunits, even across the Hfq ring (Figs. S4, 3*A*). To characterize this behavior, we mapped the interactions of individual CTDs with each of the subunits (core and CTDs) for all the models at different periods of the simulation: (i) after a 100 ns equilibration (100-110 ns, for all models), (ii) at the end of the simulation (490-500 ns for models top 1-5, 190-200 ns for models top 6-10), and (iii) over the entire 400 ns simulation (100-500 ns for models top 1-5) (Figs. S4, S5 and S6). We found that the CTDs interacted with their own core and adjacent subunits, but some CTDs also engaged with as many as 5 other subunits and CTDs. These contacts were not static, with multiple, interconnected interactions observed at different times.

We next asked how common it was for the CTDs to engage in these long-range interactions (Fig. S6). In all models, a CTD interacted with 2–6 subunit cores over a 10 ns timescale, and this degree of connectivity peaked at ~3 CTDs per hexamer (Figs. 3*B* and S6*A*). This distribution was similar at the beginning and end of the simulations, indicating convergence. When averaged over the entire 400 ns simulation, the distribution shifted to larger numbers of subunits (Fig. 3*B*, purple), suggesting that the CTDs sample different regions of Hfq over time. The CTDs also interacted amongst themselves, with a peak of three CTDs contacting other CTDs when averaged over 10 ns periods at the beginning and the end of the simulation (Figs. 3*C* and S6*B*), and five other CTDs when considering the full 400 ns simulation (Fig. 3*C*, purple). Thus, the disordered and dynamic nature of the CTDs enables a far-ranging network of interactions connecting the whole Hfq hexamer.

### RNA binding stabilizes HfqΔCTD but destabilizes WT Hfq

Nucleic acids compete with the CTDs for binding to Hfq’s rim (21), and thus, RNA binding has the potential to alter the organization of the CTDs around Hfq. Since perturbation of the CTDs destabilizes the Hfq hexamer (Figs. 2*E* and 2*G*), the stability and fragmentation pattern of the Hfq hexamer may report on the impact of RNA binding on the protein. Although the fragmentation pattern doesn’t directly reveal the RNA-protein contacts, it provides information on how the bound RNA energetically connects the Hfq subunits.

To determine whether RNA binding perturbs the structure of Hfq, we designed a series of short RNAs that mimic the Hfq binding motifs in natural sRNAs and mRNA targets of Hfq. The designed RNAs interact with different surfaces of the Hfq hexamer (Fig. 4*A* and S7). rA_6_ and rA_18_ bind two or six subunits on the distal face of Hfq (30). rU_6_ and rAU_5_G mimic the sRNA 3’ end and contact 5-6 subunits around the proximal inner surface (31–33). rCU_2_C_2_ and rim-SL, which contains rCU_2_C_2_ plus a stable stem loop, mimic RNA motifs that bind near a subunit interface within the hexamer, and interact with the arginine patch on the rim (12, 20). Additionally, we studied two larger RNAs that interact with both the proximal face and rim: rim SL-U_6_, designed to mimic an entire sRNA 3’ Hfq binding region, and RybB, a 79 nt natural sRNA that contains both the 3’ Hfq binding region and a 5’ seed region responsible for targeting the mRNA (34, 35).

**Fig. 4.**
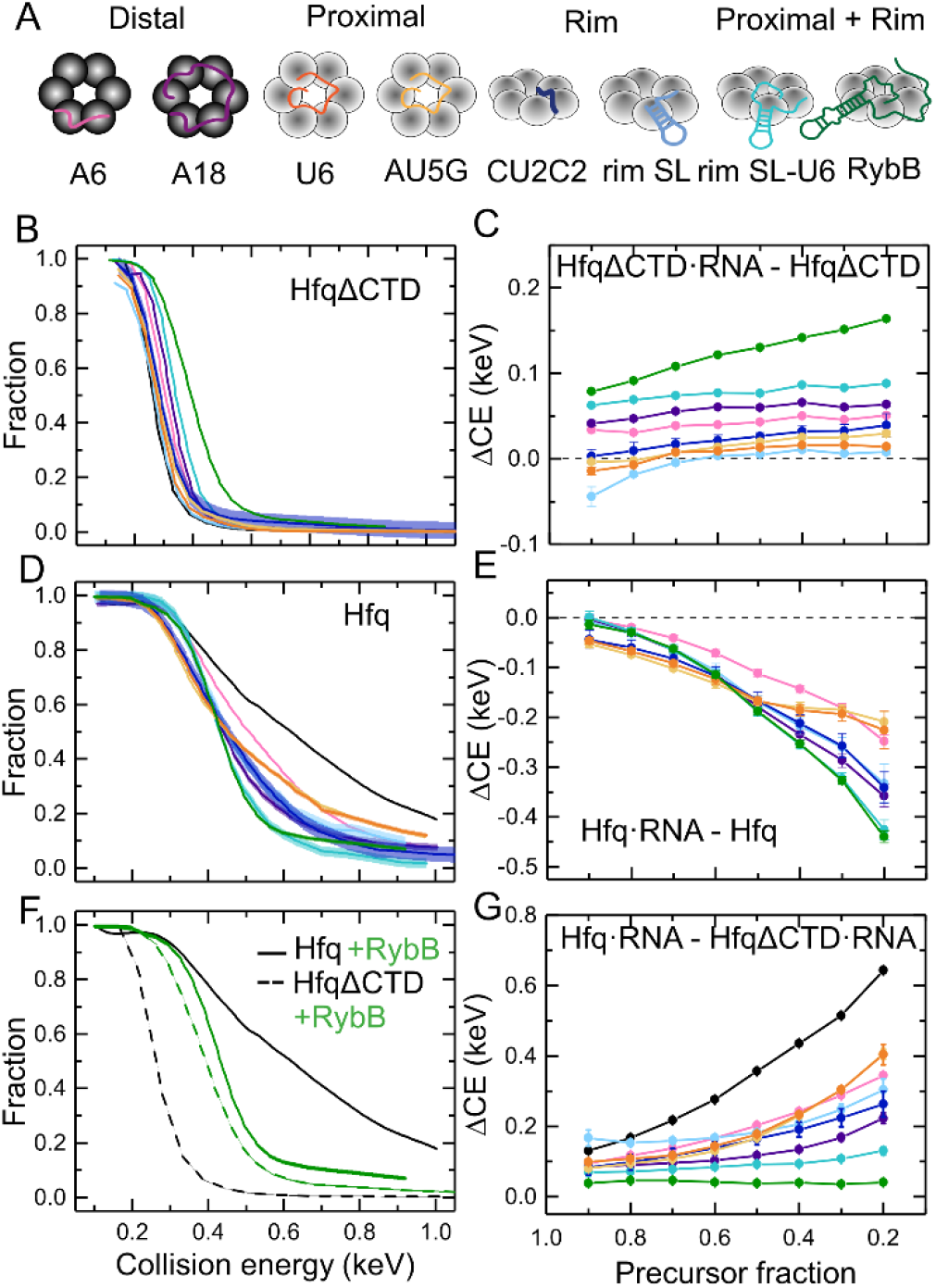
RNA binding stabilizes HfqΔCTD but destabilizes the WT protein. (*A*) RNAs used in this study (Table S2) interact with different surfaces of Hfq. (*B, D*) ERMS showing the fraction of precursor for (*B*) HfqΔCTD and (*D*) Hfq bound to RNA. Colors correspond to the RNAs in (*A*). The collision energies are corrected for the mass of the RNAs (m_Protein_/m_Protein•RNA_). The collision energies for Hfq are also corrected for the mass of the CTDs (m_HfqΔCTD_/m_Hfq_). Solid lines represent a cubic interpolation of the data. The spread on the interpolated line represents the mean of the errors of individual data points. For clarity, data symbols and error bars are not shown. See Figs. S8-S9 for further data on all dissociation products. (*F*) ERMS of Hfq (solid lines) and HfqΔCTD (dashed lines) with (green) and without (black) bound RybB. (*C, E, G*) Collision energy difference (ΔCE) between (*C*) HfqΔCTD•RNA – HfqΔCTD, (*E*) Hfq•RNA – Hfq, and (*G*) Hfq•RNA – HfqΔCTD•RNA as a function of precursor fraction. Colors are as in (*A*). Black line depicts Hfq – HfqΔCTD. Solid lines are a visual guide. Errors were calculated from the spread of the ERMS, as explained in SI Materials and Methods.

We exploited nMS to monitor the dissociation of 1:1 RNA:Hfq complexes. The mass-corrected ERMS plots of HfqΔCTD revealed a greater dissociation energy when the protein was bound to RNA, indicating that in the absence of the CTDs, the RNA stabilizes the Hfq hexamer (Figs. 4*B* and S8). To determine how much stability HfqΔCTD gained from its interactions with the RNA, we calculated the difference in collision energies between HfqΔCTD•RNA and HfqΔCTD at each remaining fraction of precursor (Fig. 4*C*). We found that all RNAs stabilized HfqΔCTD over a range of precursor fractions. Short RNAs contacting the proximal face (U_6_ and AU_5_G) and the rim (CU_2_C_2_ and rim-SL) provided minimal extra stability. The longest RNAs tested (RybB and rim-SL-U_6_) provided the most extra stability, followed by RNAs that bind to the distal face (rA_18_ and rA_6_).

Unlike HfqΔCTD, RNA binding destabilized WT Hfq, relative to free Hfq. Not only did the RNA complexes fragment at lower collision energies than the unbound protein (Figs. 4*D* and S9), but the shapes of the ERMS plots differed for the various RNAs, suggesting that the complexes have different structures. Surprisingly, the difference in collision energies between Hfq•RNA and Hfq showed that even short RNAs and RNAs that bind the distal face destabilized the complexes substantially (Fig.4*E*). Thus, in the presence of the CTDs, RNA binding can result in net destabilization of Hfq complexes.

### RNA binding removes stabilizing CTD interactions

We hypothesized that RNA binding destabilizes Hfq by perturbing the organization of the disordered CTDs. To determine how much the CTDs stabilize the complexes, we calculated the difference in collision energy needed to fragment free WT Hfq and HfqΔCTD with and without RNA (Figs. 4*F-G*). In the absence of RNA, the CTDs significantly stabilized the Hfq hexamer (Fig. 4*F-G*, black lines). In the presence of RybB sRNA, the complexes had similar stabilities that were intermediate between apo WT Hfq and apo HfqΔCTD (Fig. 4*F*, green lines). This result suggested that RNA binding opposes the effects of the CTDs while stabilizing the Hfq core.

For the free proteins, the observed stability gap was wider as more precursor was fragmented (Fig. 4*G*), suggesting a range of complexes with variable strengths of CTD interactions, as already deduced by comparison of the protein’s ERMS (Figs. 2*B-C*). Upon RNA binding, the stability conferred by the CTDs became more uniform as the precursor was dissociated (Fig. 4*G*, colored lines). Thus, RNA binding results in complexes that dissociate more homogeneously.

All the RNAs tested interfered with stabilization by the CTDs. The effect was strongest for the longest RNAs (rim SL-U_6_ and RybB) that presumably displace more CTD contacts with the rim and proximal face of Hfq. However, even the smallest RNAs tested abrogated the stabilizing effect of the CTDs (Fig. 4*G*). Interestingly, this loss was also substantial for RNAs binding the distal face, even though only the first few residues of the CTDs are close to this surface (10).

Next, we studied how RNA binding affected the dissociation pathways of Hfq and HfqΔCTD, by determining which subcomplexes retained the RNA after hexamer dissociation (Fig. S10*A*). To minimize contributions from secondary dissociation events, we compared the dissociation products when only 20% of the protein was fragmented (0.8 precursor fraction). Comparison of the dissociation pathways revealed similar fragmentation of RNA complexes with Hfq and with HfqΔCTD (Fig. S10*B-C*). This result suggested that in WT Hfq, RNA binds to the core and that perturbation of the CTDs likely comes from their displacement from the core and not from direct interactions with the RNA. CTD displacement was maximal for the largest RNA (RybB sRNA), as the ERMS for Hfq and HfqΔCTD almost overlapped (Fig. 4*F*). The ERMS of shorter RNA complexes were closer to the ERMS of the free proteins, indicating that binding to the core did not require as much displacement of the CTDs (Figs. 4*B* and 4*D*). Finally, we observed that the percentage of fragments with bound rim SL was different for each protein, suggesting different modes of binding that could explain the lesser perturbation of the CTDs by this RNA (Figs. S10*C* and 4*G*; light blue). The dissociation of Hfq•AU_5_G was also slightly altered, in contrast with U_6_’s, which was identical with or without the CTDs (Fig. S10*C*; orange and yellow). Given that the fragmentation of both proximal-binding RNAs was very similar for the truncated protein, it is possible that the CTDs help discriminate between optimal proximal-binding RNA motifs.

### Progressive RNA binding displaces the CTDs from Hfq

The design of our RNAs allowed us to investigate the interplay between Hfq, the disordered CTDs and RNA-protein interactions as an sRNA progressively binds to the protein (Figs. 5*A-B* and S7*B*). For this, we compared the stabilities and dissociation pathways of WT Hfq and HfqΔCTD in the absence and presence of RNA segments mimicking a stepwise binding process (Fig. 5*B*). On the one hand, we found that as the RNA interacts with more surfaces of Hfq, the RNA conferred stability to the protein core (Fig. 5*C*; HfqΔCTD). However, RNA binding was accompanied by a loss of favorable CTD interactions (Fig. 5*C*; CTD) that reduced stability overall (Fig. 5*C*; Hfq). On the other hand, progressive RNA binding shifted the dissociation products to increasingly larger fragments (Fig. 5*D*). This dissociation pattern was the same for Hfq and HfqΔCTD, indicating similar RNA interactions with the core in both proteins. Thus, stable interactions between RNA and the core are established as the CTDs are displaced.

**Fig. 5.**
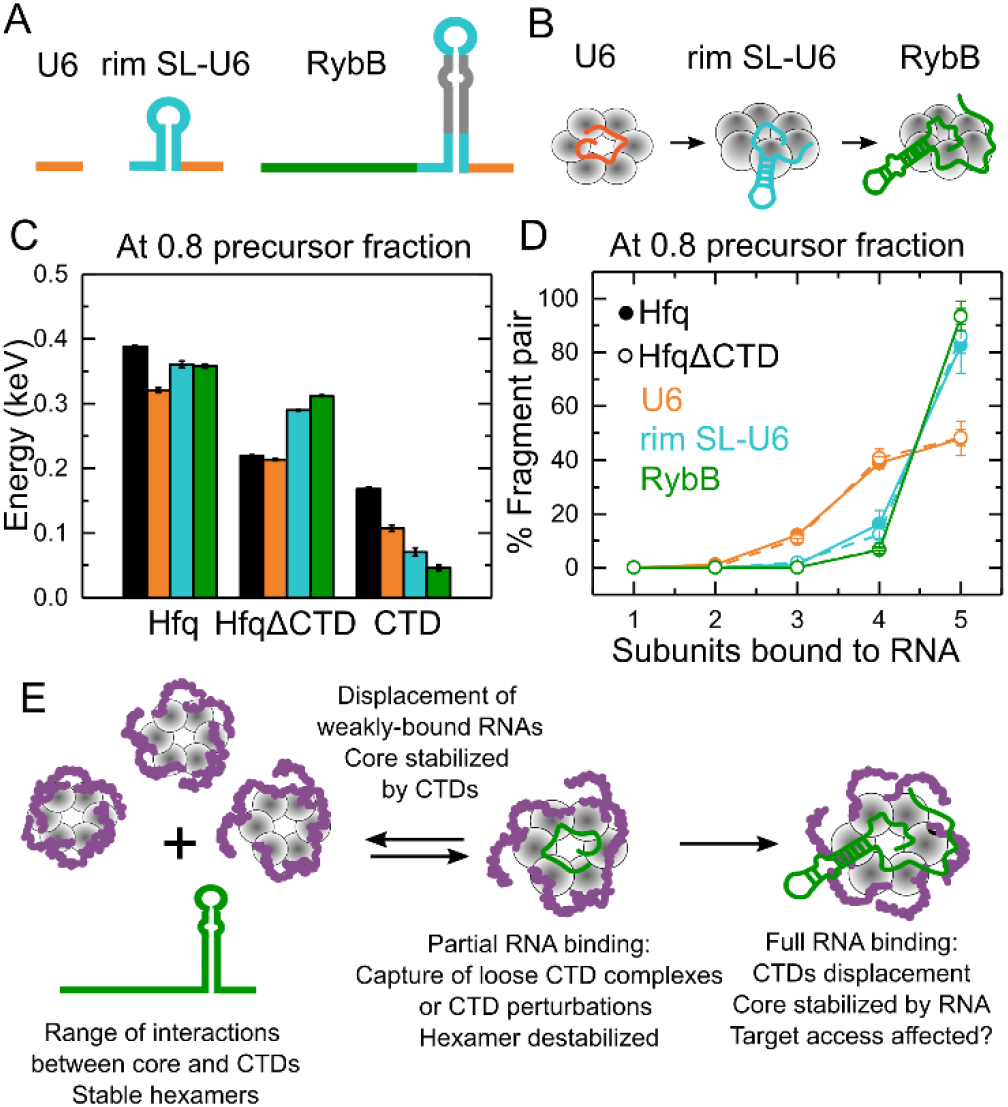
RNA binding progressively displaces the Hfq CTDs. (*A*) RNAs mimicking segments of RybB sRNA that progressively interact with more Hfq surface: 3’ U tail (orange), terminal stem-loop (cyan), 5’ seed (dark green). (*B*) Cartoon of progressive RNA binding. (*C*) Stabilities of Hfq, HfqΔCTD (core) and CTDs when bound to the RNAs shown in (*A*). Colors correspond to the RNAs as shown in (*B*). The stabilities were determined as the collision energies of Hfq, HfqΔCTD or their difference when 20% of the precursor was fragmented (0.8 precursor fraction). Errors were calculated as in Fig. 4*C* (see SI Methods). (*D*) Percentage of fragment pairs after the collision of Hfq (solid lines) or HfqΔCTD (dashed lines) bound to the RNAs in (*A*) (see also Fig. S10). Errors were calculated as for Figs. 2*D-E*. (*E*) Assorted organization of the disordered CTDs results in heterogeneous complexes. RNAs could bind complexes with loose CTD conformations or perturb a CTD. Local CTD disturbances destabilize the hexamer and are transmitted to the rest of the protein, promoting further CTD displacement and stable binding for favorable RNAs. Non-preferred RNAs are displaced by competition with the CTDs (21), reestablishing hexamer stability

## Discussion

In this work, we employed nMS-SID to analyze Hfq complexes of defined stoichiometry, connectivity and shape. The results demonstrate that the CTDs connect all of the Hfq subunits, stabilizing the hexamer (Fig. 2*C*, 2*G* and S10*A*). Moreover, MD simulations using an improved force-field for IDPs (36) revealed that each CTD can dynamically interact with several other CTDs and folded domains within the hexamer (Figs. 3, S3 and S6). Although RNA binding also stabilizes the folded core of Hfq, RNA disrupts the network of CTD interactions. Only RNAs that form multiple favorable contacts with Hfq offset the loss of stabilization by the CTDs, explaining how the CTDs help Hfq discriminate among different RNAs.

Based on our results, we propose that RNAs stably bind Hfq through a stepwise process (Fig. 5*E*). An RNA may first take advantage of configurations in which an Hfq region is exposed, or a CTD is loosely folded. The initial interaction with a segment of the RNA perturbs nearby CTDs. Because the CTDs are interconnected, this perturbation propagates to the rest of the protein hexamer, resulting in more CTD displacement and further RNA binding. If binding is favorable, the hexamer is stabilized. However, if binding is not favorable, the hexamer is destabilized, making it more favorable for the CTDs to regain their interactions with the rest of the protein. This search for stability could explain the removal of weakly-bound RNAs (21). Additionally, an integrated organization of the CTDs could facilitate sRNA competition and the search for mRNA targets, as binding of a second RNA will also perturb the CTDs, stimulating sRNA displacement or disassembly of non-cognate ternary complexes. Finally, the varied CTD conformations impart asymmetry to the Hfq homohexamer, that may also contribute to the selection of RNA ligands.

Our nMS-SID results showed that the CTDs make the Hfq hexamer more resistant to dissociation by collision (Fig. 2*C*). This observation agrees with previous collision-induced dissociation MS experiments showing that *E. coli* Hfq is more stable than Hfq from *V. cholerae*, which has a 22 residue CTD (37). Vincent et al. (37) attributed this stabilization to packing of the CTDs along the intersubunit interfaces. Based on our observation that the CTDs unfold before the subunits dissociate, we propose that Hfq is additionally stabilized by CTD interactions with multiple Hfq subunits that are facilitated by longer CTDs.

Network connectivity can explain why the binding of even short RNAs reduced the CTD’ s stabilizing contribution (Figs. 4*D-G* and 5*C*). Also, since RNA-bound HfqΔCTD and Hfq fragmented similarly (Figs. 5*D* and S10), stable RNA binding seems to involve contacts with the core and not the CTDs, as previously proposed (12, 38, 39). Finally, destabilization of Hfq but not HfqΔCTD when A18 RNA is bound (Fig. 4*C*) supports a reported functional link between the distal RNA binding face and R66 at the start of the CTD (23).

Hfq is a model protein with disordered regions that act synergistically to communicate perturbations among its subunits. This feature is enabled by an architecture of identical monomers, each providing an identical disordered region. The single-stranded DNA binding (SSB) protein, a homotetramer involved in DNA repair, replication and recombination (40), also contains disordered CTDs with parallels to Hfq: they impart stability to the SSB tetramer (41) but are displaced upon partial DNA engagement, modulating binding and SSB oligomerization (42). Importantly, partial deletion of SSB’s CTDs results in impaired activity, indicating a critical role for multiple CTD interactions in cellular function (43). Histone tails are also thought to fold heterogeneously around the DNA (44, 45), and to be disrupted by chaperone binding or post-translational modification (46). It would be interesting to know if other RNA-binding proteins use interconnected IDPs to integrate the molecular interactions within RNA-protein complexes.

## Materials and Methods

### RNA preparation

RybB RNA was transcribed using phage T7 RNA polymerase followed by 8% polyacrylamide gel purification (8M urea). The remaining short RNAs used in the study were purchased from IDT with HPLC purification. See Table S2 for RNA sequences.

### Protein expression and purification

HfqΔCTD and Hfq were purified as described before (20).

### Sample preparation for native mass spectrometry (nMS)

Protein (HfqΔCTD and Hfq) and RNA (rA_18_, rim SL-U_6_ and RybB) samples were dialyzed overnight into 500 mM ammonium acetate pH 6.8 (99.99%, MilliporeSigma), with eight buffer exchanges (3.5 kDa mass cutoff micro-dialysis, Pierce). This ionic strength prevented precipitation of protein-RNA complexes at the concentrations (10 μM hexamer) required for ion mobility MS. The remaining RNA samples were used as supplied by the manufacturer and did not require additional desalting. Protein-RNA complexes were prepared by mixing 1:1 RNA and HfqΔCTD or Hfq to a final concentration of 10 μM each. 400 mM ammonium acetate (final concentration) plus triethylammonium acetate (TEAA) (1 M, MilliporeSigma; 100 mM final concentration for charge reduction) were subsequently added to the samples.

### Mass spectrometry

All samples were introduced into the mass spectrometer using nano-electrospray emitters that were prepared in-house using a Sutter P-97 micropipette puller. All spectra in this work were acquired on a Waters Synapt G2 HDMS instrument (Waters Corporation, Wilmslow, U.K.) modified with a surface-induced dissociated (SID) device between a shortened trap stacked ring ion guide and an ion mobility cell, as described previously (47). SID lenses can be tuned either to transmit ions for MS or to direct the ions onto the surface for collision. Typical voltage settings and instrument parameters used here for transmission mode and SID can be found in the Supporting Information (Tables S3-S4). Energy resolved mass spectra (ERMS) were produced by acquiring data from tandem MS experiments with SID voltage potentials ranging from 15 and 140 V. Each experiment was repeated in technical triplicate. Additional information is provided in SI.

### Analysis of mass spectrometry data

Ion mobility was used to separate product ions and selection rules for each SID product were made using Waters Corporation Driftscope 2.9 software. The intensity of subcomplexes were extracted from SID spectra with TWIMExtract v1.3 (48). Collision energies were calculated as *E*(*eV*) = *zV_SID_;* where *z* is the charge state of the precursor ions and *V_SID_* is the SID voltage. Energy-resolved mass spectra (ERMS) were corrected by the *m_HfqΔCTD_/m_Hfq_* and *m_Protein_/m_Protein–RNA_*. Additional information provided in SI.

### MD simulations

All simulations were performed with the molecular dynamics program OpenMM (49) and CHARMM36m force-field (50). Simulations were started from the PDB ID 1HK9 which included residues 7-68 (29). The starting structures of the missing CTDs were obtained from a) top 10 Rosetta models (15) (top 1-10) and b) 1 structure in which CTDs are fully extended (21). Four simulations with extended CTDs were performed starting with different initial velocities. All protein structures were embedded in a water box and neutralized with 150 mM Na^+^Cl^-^. Additional simulation and setup details are provided in the SI. Following a 100 ns equilibration, the models were run for an additional 400 ns (top 1-5), 100 ns (top 6-10) or 10 ns (extended 1-4) (Table S5). To gain information about the short-term structures and their evolution, we analyzed contacts between CTDs and cores of various subunits during various time intervals of the simulation (Figs. S4-S6). Additional information provided in SI.

## Acknowledgements

The authors thank Benjamin Jones for his assistance with instrument maintenance and automated data acquisition and Andrew Santiago-Frangos for help with initial samples. This work was funded by grants from National Institutes of Health [1P41GM128577-01 to V.H.W.; R35 GM136351-03 to S.A.W.], the Johns Hopkins University Provost’s Office [PURA to H.L.]. Part of this research was conducted using computational resources at the Maryland Advanced Research Computing Center.

