## Supplementary Information for "Intrinsically disordered interaction network in an RNA chaperone revealed by native mass spectrometry"

### **Supplementary Information for**
Intrinsically disordered interaction network in an RNA chaperone revealed by native mass spectrometry.

Samantha H. Sarni[1†] (1), Jorjethe Roca[2] (1), Chen Du[1], Mengxuan Jia[1‡], Hantian Li[2], Ana Damjanovic[2], Ewelina M. Małecka[2], Vicki H. Wysocki[1*], Sarah A. Woodson[2*]

Vicki H. Wysocki or Sarah A. Woodson


**This PDF file includes:**

Supplementary text
Figures S1 to S10
Tables S1 to S5
SI References

#### Supplementary Information Text

#### SI Materials and Methods

**Mass spectrometry.** Nano-electrospray ionization (nESI) emitters were prepared in-house using a Sutter P-97 micropipette puller. 3-6 µL samples were loaded into the nESI emitters using a Hamilton 10 µL syringe or ultra-micro gel loading pipette tips. All spectra in this work were acquired on a Waters Synapt G2 HDMS instrument (Waters Corporation, Wilmslow, U.K.) modified with a surface-induced dissociation (SID) device, as described previously (1). Briefly, a custom SID device was inserted between a shortened trap stacked ring ion guide and an ion mobility cell. Voltages were supplied to the SID cell via external DC power supplies (Ardara Technologies, Ardara, PA) and controlled through the accompanying Tempus Tune software (Ardara Technologies, Ardara, PA). SID lenses can be tuned either to transmit ions for MS or to direct the ions onto the surface for collision. Typical settings used here for transmission mode and SID can be found in Tables S3-S4. Energy resolved mass spectra (ERMS) were produced by acquiring data from tandem MS experiments with SID voltage potentials ranging from 15 and 140 V. The precursor charge state for SID experiments was chosen as the lowest available charge state (which retains native-like state more reliably) that has a stable enough signal intensity for 21 different SID spectra. Each experiment was repeated in technical triplicate.

**Analysis of mass spectrometry data.** The expected molecular masses of monomer Hfq and RNAs were calculated using the UniDec Protein/RNA mass calculator (1). Selection rules, arrival time, and *m/z* were made for each SID product using Waters Corporation Driftscope 2.9 software. The intensity of each subcomplex respective charge state was then extracted from each series of SID spectra using TWIMExtract v1.3 (2). The intensities of all charge states of each SID product were summed and averaged for three replicates and plotted with standard error. Due to the sparse nature of the mass spectra, the ERMS data underwent a cubic interpolation processing step to create the continuous data needed for comparing differences in dissociation energies. The mean of the error of data points in individual ERMS was assigned as the error of the interpolated data. Collision energies were calculated as $E(eV) = zV_{SID}$; where $z$ is the charge state of the precursor ions and $V_{SID}$ is the SID voltage defined as the potential difference between the trap exit and the surface. To eliminate the mass difference between HfqΔCTD and Hfq, the collision energies were corrected by the factor $m_{Hfq\Delta CTD}/m_{Hfq}$ (3). Collision energies for protein-RNA complexes were further corrected by the factor $m_{Protein}/m_{Protein-RNA}$ to account for the mass provided by the bound RNA.

Surface-induced unfolding (SIU) plots were generated by extracting and normalizing the intensity of the respective precursor ion using TWIMExtract and ORIGAMI 1.2.1.6 (4).

Errors in collision energy difference (ΔCE) plots (Figs. 4*C, E,* and *G*) were propagated as $\sigma(\Delta CE) = \sqrt{\sigma^2(CE_{Protein-RNA}) + \sigma^2(CE_{Protein})}$; where $\sigma^2$ is the spread of the ERMS interpolated data on the collision energy axis (Figs. 4*B* and *4D*). Errors for Fig. 4*G*, were calculated similarly but for Hfq•RNA - HfqΔCTD•RNA.

**Collisional cross section calculation.** The experimental collisional cross sections (CCSs) were measured in a travelling wave ion mobility (TWIM) drift cell filled with $N_2$ gas. C-reactive protein, transthyretin, β-lactoglobulin A, alcohol dehydrogenase, concanavalin A and avidin were purchased from MilliporeSigma and used as calibrants. HfqΔCTD, Hfq and calibrants were buffer exchanged and charged-reduced by triethylamine acetate as described previously. Theoretical CCSs of calibrants were obtained from a database of charge-reduced proteins (5). The calculated CCSs of analytes were simulated using projected superposition approximation (PSA) on the webserver: http://psa.chem.fsu.edu/ (6). The PDB structure 1hk9 was used as a model for HfqΔCTD (7). Model top 1 from the MD simulations in this work (Fig S3) and previously published Rosetta structures (8) were used as models for Hfq.

**MD simulations.** All simulations were performed with the molecular dynamics program OpenMM (9). The top 10 models of full-length *E. coli* Hfq previously obtained with Rosetta FloppyTail (8) (models top 1-10) were used as starting structures for the MD simulations. CHARMM-GUI (10) was used for system setup – the structures were embedded in a water box with an additional 15 Å layer of water in each direction, and charge neutralized with 150 mM $Na^+Cl^-$. Additional 4 simulations (extended models 1-4) were performed by starting from an extended model of the CTDs. The initial extended model was created by prepending or appending N and C-terminal residues in a beta conformation (8) to the core of *E. coli* Hfq (1HK9; (7)). A 10 Å layer of water was added in each direction, and 150 mM Na+$Cl^-$. The protein structures quickly collapsed, so the effective water layer in the simulations was larger. The simulations were performed in an NpT ensemble, using the particle mesh Ewald method for electrostatic interactions with a real space interaction cutoff of 12 Å. We used CHARMM36m, an improved force-field for folded and intrinsically disordered proteins (11). The first 100 ns were used to equilibrate the systems and were not included in the analysis of the simulation results. Models top 1-5 were run for an additional 400 ns, models top 6-10 were run for an additional 100 ns and extended models were run for an additional 10 ns. The simulation parameters are summarized in Table S5. To gain information about the structures and their evolution, we analyzed contacts between the CTDs and the cores of the Hfq subunits during various intervals of the simulation (Figs. S4-S6). A CTD of a subunit was considered to contact the core (res 1-65) or CTD (res 66-102) of another subunit if, during the time interval of consideration, the distance between any atom of that CTD and any atom of the core (or another CTD) was less than 4 Å.

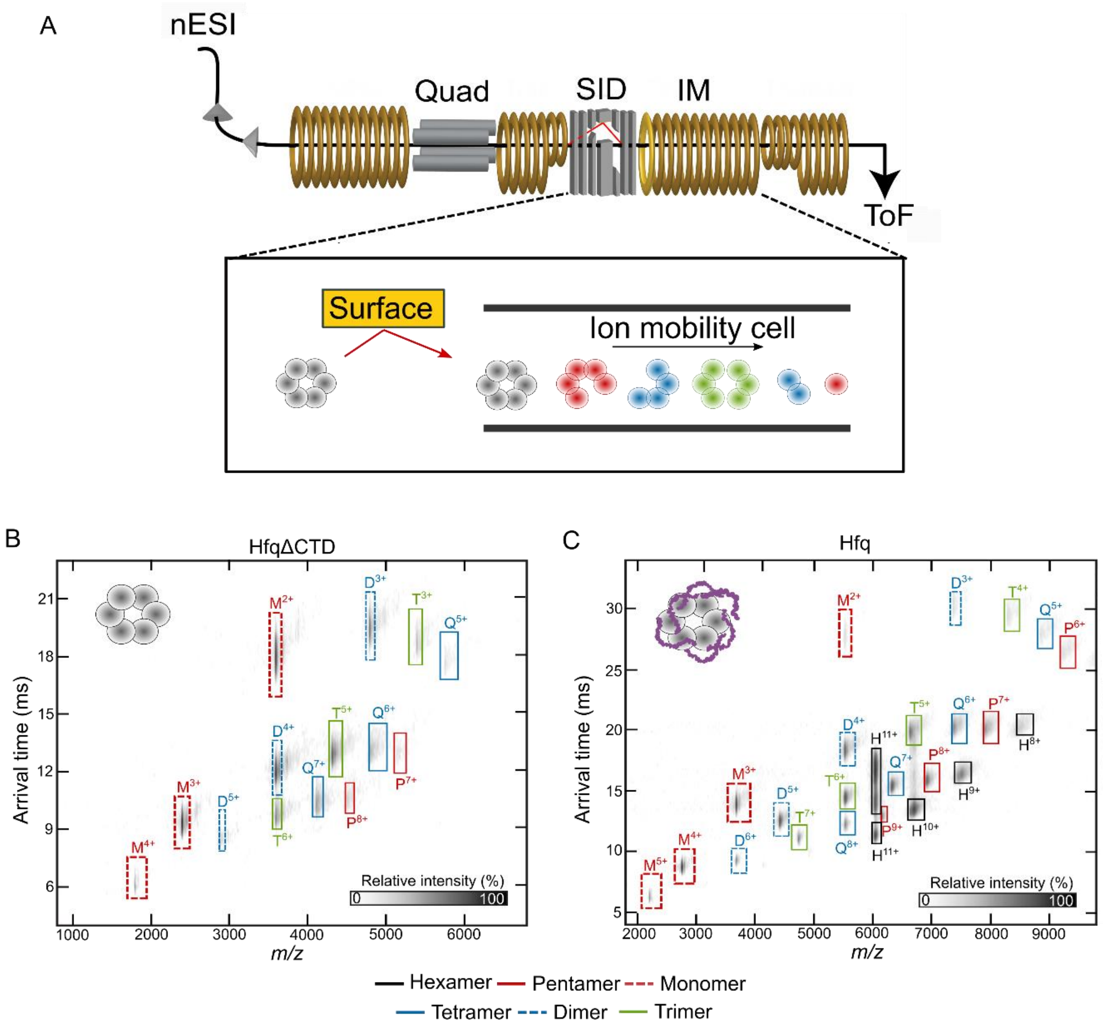


**Fig. S1.** Separation of dissociated complexes by ion mobility. (*A*) Schematic of native mass spectrometer with surface-induced dissociation (nMS-SID). Intact complexes undergo nano electrospray ionization (nESI) followed by selection of a precursor charge state with a quadrupole mass filter (Quad). The selected complex collides with a gold surface with a self-assembled monolayer (SID; red path). The resulting fragments along with the remaining precursor are separated based on mass, shape and charge in an ion mobility cell (IM) and analyzed in a time-of-flight analyzer (ToF) with results reported in an ion mobiligram. The black path indicates the trajectory of the complexes when the instrument is in transmission mode (no collisions). (*B*-*C*) Ion mobiligrams of (*B*) HfqΔCTD at SID 605 eV and (*C*) Hfq at SID 608 eV. The mobiligrams report the time needed to traverse an ion mobility cell and reach the detector (arrival time) versus the mass-to-charge ratio of the various fragments. The complexes overlap in mass-to-charge but are characterized by unique arrival times, signaling different oligomeric states: pentamer, tetramer, trimer, dimer, and monomer. Cases in which a given fragment (for example, hexamer) has the same *m/z* but different arrival times represent different conformations, with the slower drift characteristic of more extended conformations.

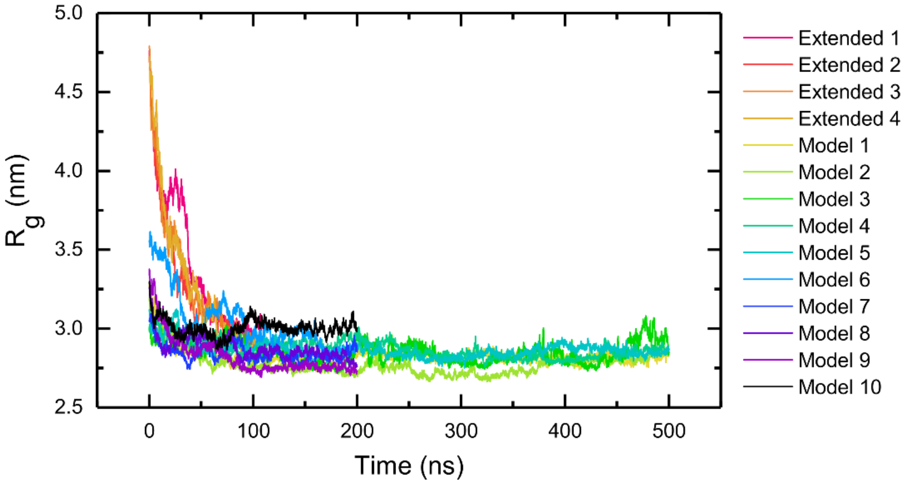


**Fig. S2.** Hfq compaction from MD simulations. Time evolution of the radius of gyration ($R_g$) of Hfq. Extended models 1-4 were run for 110 ns, models top 1-5 for 500 ns and models top 6-10 for 200 ns.

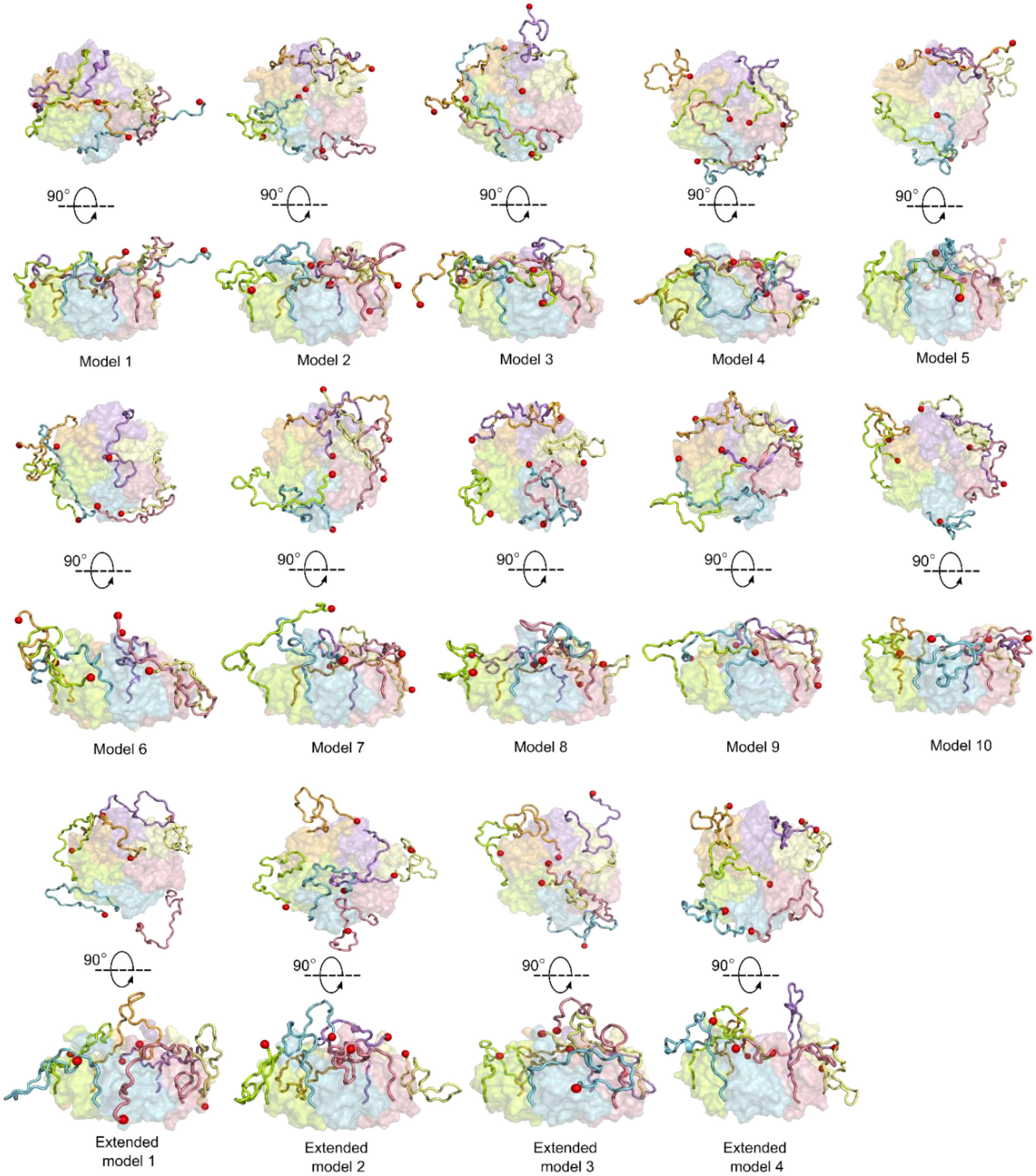


**Fig. S3.** Structures of *E. coli* Hfq from MD simulations. Top and side views of Hfq models from 14 MD runs. Monomers and their CTDs are colored individually with each acidic tip shown as a red sphere. The structures shown are from the last frame of each simulation: 500 ns for models top 1-5, 200 ns for models top 6-10 and 110 ns for extended models 1-4.

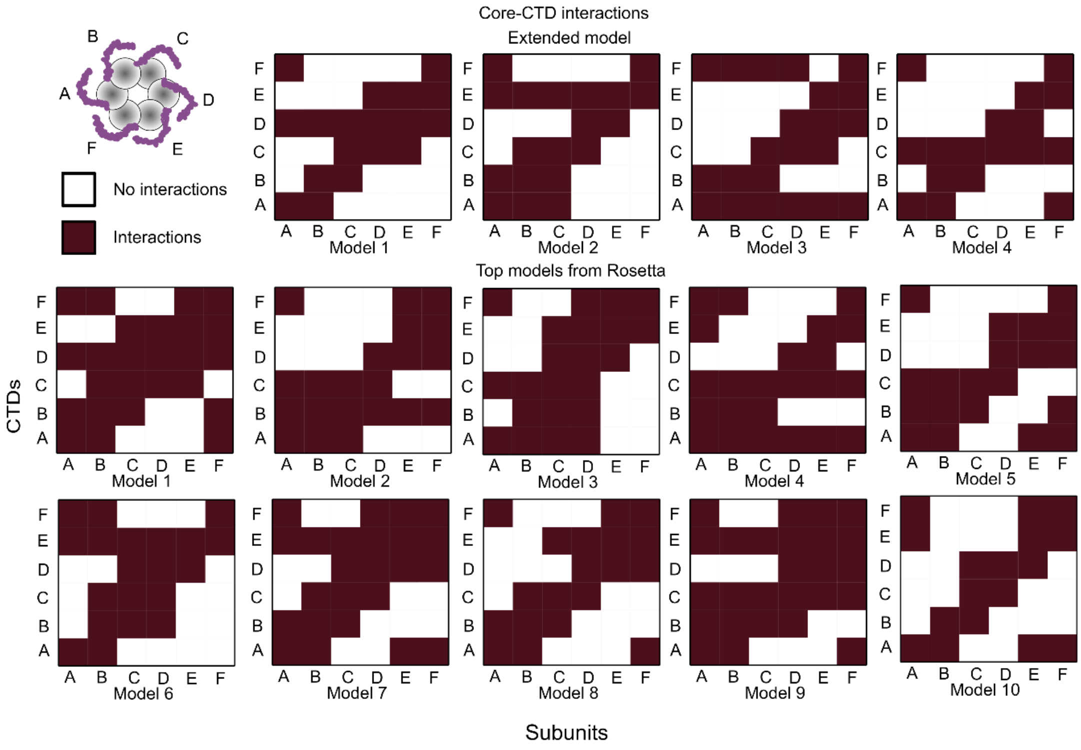


**Fig. S4.** Interactions between CTDs and subunits of Hfq. Colors in the heat map indicate that at least one residue in the CTD interacts with one residue of the core in any frame of the first 100-110 ns of the simulations, for all models; this represents the period after equilibration of the trajectories. The 6 individual subunits are labeled A-F, as illustrated in the cartoon.

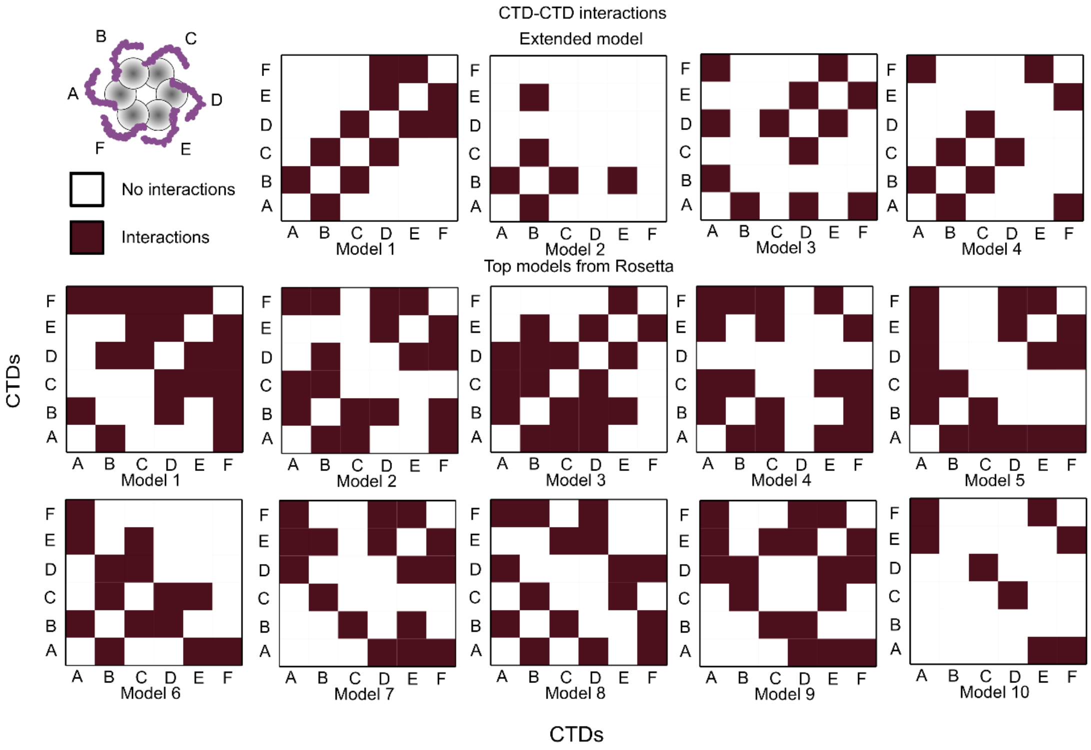


**Fig. S5.** Interactions between CTDs of Hfq. Colors in the heat map indicate that at least one residue in the CTD interacts with one residue of another CTD in any frame of the first 100-110 ns of the simulations, for all models; this represents the period after equilibration of the trajectories. The 6 individual subunits are labeled A-F, as illustrated in the cartoon.

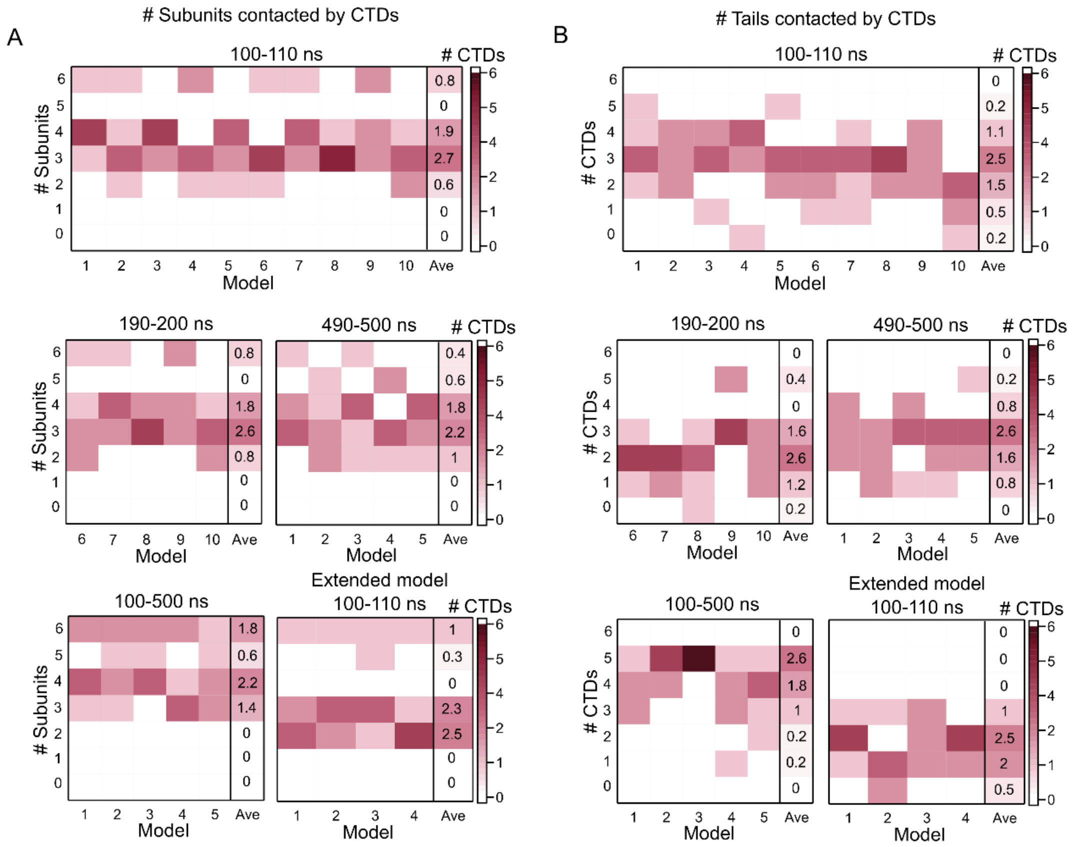


**Fig. S6.** Hfq compaction from MD simulations. Disordered CTDs establish multiple interactions across Hfq. (*A*-*B*) Number of CTDs interacting with 1, 2, 3, 4, 5, or 6 (*A*) subunits or (*B*) CTDs of Hfq for the different models obtained by MD simulations.

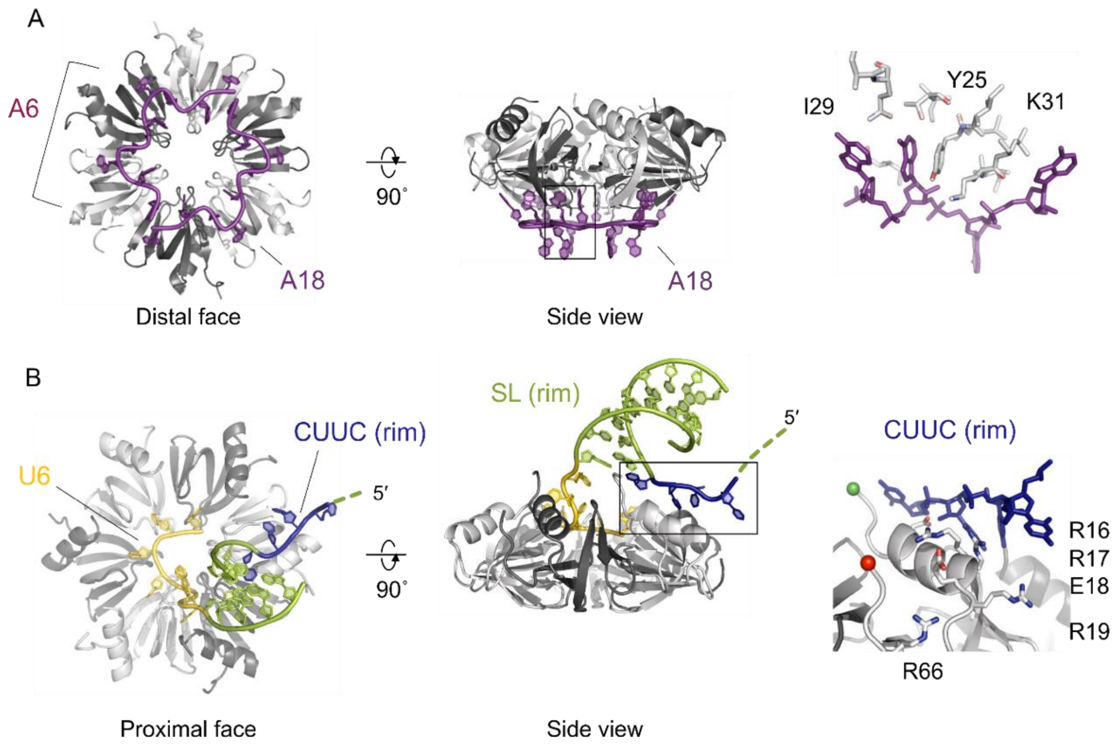


**Fig. S7.** RNA interactions with Hfq. (*A*) $rA_{18}$ bound to the distal face with three nucleotides per Hfq subunit, from 3GIB (12). $rA_6$ is expected to only bind two subunits. (*B*) RydC sRNA bound to the proximal face and rim, from 4V2S (13). The 3′ terminal uridines (gold) bind around the proximal pore, the terminal stem-loop (SL; green) sits atop the proximal face, and the single-stranded CUUC motif (blue) interacts with the arginine patch on the rim of the hexamer. RybB sRNA and the minimal rim-SL RNA contain similar sequences and are expected to bind Hfq similarly. The RNAs are color-coded as in Fig. 4*A*.

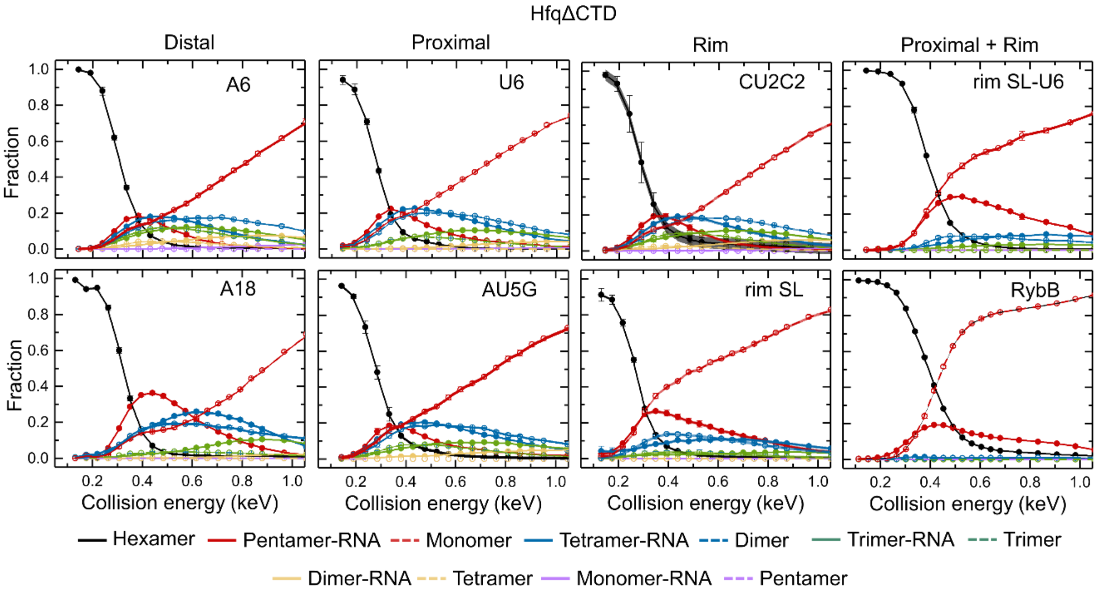


**Fig. S8.** RNA binding stabilizes Hfq's core. Energy-resolved mass spectra (ERMS) of HfqΔCTD bound to the indicated RNAs. The collision energies are corrected to account for the mass of the RNAs ($m_{Hfq\Delta CTD}/m_{Hfq\Delta CTD \bullet RNA}$). Reported fractions are the sum of the intensities of each dissociation product (from ion mobiligrams) normalized by the total intensity of all products. Symbols report the average of three replicates and the standard errors, which are smaller than symbols for some data points. Solid lines represent a cubic interpolation of the data. The spread (negligible) on the interpolated line represents the mean of the errors of individual data points.

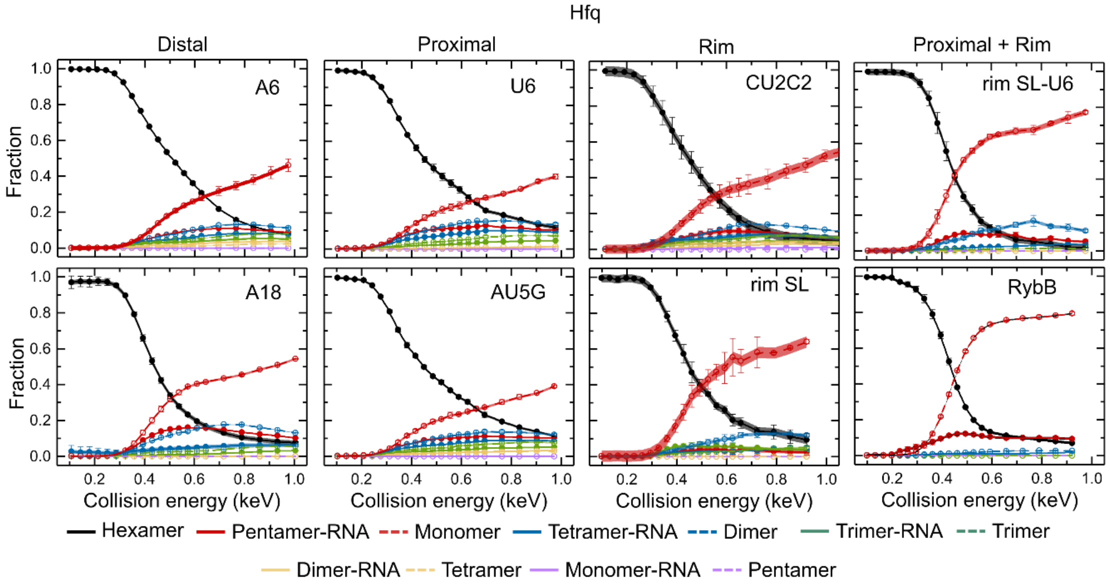


**Fig. S9.** RNA binding destabilizes Hfq. Energy-resolved mass spectra (ERMS) of Hfq bound to the indicated RNAs. The collision energies are corrected to account for the mass of the RNAs ($m_{Hfq}/m_{Hfq•RNA}$) and the CTDs ($m_{Hfq\Delta CTD}/m_{Hfq}$). Reported fractions are the sum of the intensities of each dissociation product (from ion mobiligrams) normalized by the total intensity of all products. Symbols report the average of three replicates. Errors are the standard error and are smaller than symbols for some data points. Solid lines represent a cubic interpolation of the data. The spread (negligible) on the interpolated line represents the mean of the errors of individual data points.

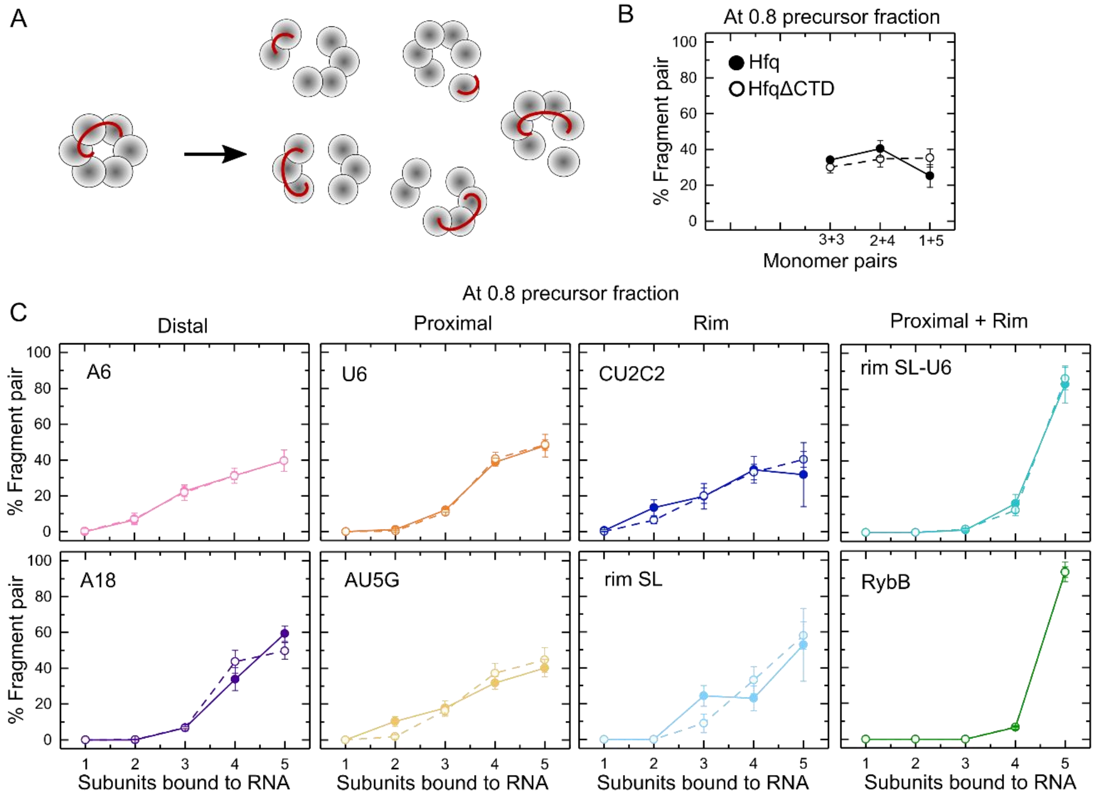


**Fig. S10.** Dissociation pathways of Hfq complexes. (*A*) Hfq bound to RNA dissociates into different pathways with different fragment pairs (pentamer•RNA + monomer, etc.). (*B*) Percentage of fragment pairs obtained after the collision of Hfq (solid lines) or HfqΔCTD (dashed lines). (*C*) Percentage of fragment pairs obtained after the collision of Hfq (solid lines) or HfqΔCTD (dashed lines) bound to the indicated RNAs. The fragments pairs were calculated for when 20% of the precursor is fragmented (0.8 precursor fraction). Errors are the addition of the spread of the ERMS curves for the fragment pairs (Figs. S8-S9), normalized by the total dissociated fraction and converted to percentage. Solid lines are a visual guide.

**Table S1.** Experimental (TWIM) and calculated (PSA) collision cross sections (CCS) of HfqΔCTD and Hfq.

| Protein | TWIM[a] | | PSA[a] | |
|---|---|---|---|---|
| | Charge state | CCS (Å$^2$)[b] | Model[c] | CCS (Å$^2$) |
| HfqΔCTD | 10+ | 3341±17 | PDB:1hk9 | 3243 |
| | 9+ | 3278±12 | | |
| Hfq | 12+ | 4125±19 | MD | 4473±129[d] |
| | 11+ | 4096±14 | Rosetta | 5539 |

[a]TWIM: Traveling wave ion mobility. PSA: Projected superposition approximation.

[b]Error of TWIM is shown as standard deviation of triplicate measurements.

[c]Models for calculated CCS: PDB from database (https://www.rcsb.org/) and Sauter et al. (7); MD from this work; Rosetta from Santiago-Frangos et al. (8).

[d]PSA-calculated CCS of Hfq is shown as average and standard deviation of models top 1-5 from MD simulations.

**Table S2.** Sequences of RNAs used in this work.

| RNA | Sequence (5'→3') |
|---|---|
| $rA_6$ | AAAAAA |
| $rA_{18}$ | AAAAAAAAAAAAAAAAAA |
| $rU_6$ | UUUUUU |
| $rAU_5G$ | AUUUUUG |
| $rCU_2C_2$ | CUUCC |
| rim SL | CUUCCGUCCAUUUCGGACG |
| rim SL-$U_6$ | CUUCCGUCCAUUUCGGACGUUUUUU |
| RybB[a] | gGCCACUGCUUUUCUUUGAUGUCCCCAUUUUGUGGAGCCCAUCAACCCCGCCAUUUCGGUUCAAGGUUGAUGGGUUUUUU |

[a]Nucleotides in lowercase letters were added to the natural RybB sequence to facilitate *in vitro* transcription by T7 polymerase.

**Table S3.** Typical voltage parameters on components of the modified Waters SYNAPT G2 mass spectrometer used in this work for surface-induced dissociation (SID).

| **Component** | **Voltage (V)** | | |
|---|---|---|---|
| | Transmission mode[a] | SID 15 (V) | SID 15+x* (V)[b] |
| Trap bias | 45 | 75 | 75+x* |
| Trap exit | -40 | -15 | -15+x* |
| Entrance 1 | -42 | -15 | -15+x* |
| Entrance 2 | -44 | -46 | -46 |
| Front top | -63 | -56 | -56 |
| Front bottom | -64 | -15 | -15+x* |
| Middle bottom | -61 | -46 | -46 |
| Surface | -59 | -30 | -30 |
| Rear top | -60 | -170 | -170 |
| Rear bottom | -61 | -55 | -55 |
| Exit 1 | -73 | -67 | -67 |
| Exit 2 | -74 | -77 | -77 |

[a]Transmission mode: The complexes are not made to collide with the soft surface, instead directly passing to the ion mobility cell (Fig. S1*A*).

[b]x*: SID voltage increment compared to SID 15V. To produce higher SID voltages, the voltages applied on the trap cell and two SID lens were increased.

**Table S4.** Parameters used for nMS-SID experiments.

| **Parameter** | **Value** |
| --- | --- |
| Sampling cone voltage | 20 V |
| Extraction cone voltage | 2 V |
| Source temperature | 30 °C |
| Trap gas flow | 2 mL/min |
| Ion mobility gas flow | 60 mL/min |
| Ion mobility wave velocity | 320 m/s |
| Ion mobility wave height | 20 V |

**Table S5.** Molecular dynamics simulation parameters.

| **Model** | **Starting model**[a] | **Length of simulation (ns)** |
|---|---|---|
| top1 | Rosetta top 1 | 500 |
| top2 | Rosetta top 2 | 500 |
| top3 | Rosetta top 3 | 500 |
| top4 | Rosetta top 4 | 500 |
| top5 | Rosetta top 5 | 500 |
| top6 | Rosetta top 6 | 200 |
| top7 | Rosetta top 7 | 200 |
| top8 | Rosetta top 8 | 200 |
| top9 | Rosetta top 9 | 200 |
| top10 | Rosetta top 10 | 200 |
| Extended 1 | Hfq with extended CTDs 1 | 110 |
| Extended 2 | Hfq with extended CTDs 2 | 110 |
| Extended 3 | Hfq with extended CTDs 3 | 110 |
| Extended 4 | Hfq with extended CTDs 4 | 110 |

[a]Starting Rosetta models correspond to structures with the lowest free energies as produced in (8).
